# Huntingtin confers fitness but is not embryonically essential in zebrafish development

**DOI:** 10.1101/615591

**Authors:** Harwin Sidik, Christy J. Ang, Mahmoud A. Pouladi

## Abstract

Attempts to constitutively knockout HTT in rodents resulted in embryonic lethality, curtailing efforts to study HTT function later in development. Here we show that HTT is dispensable for early zebrafish development, contrasting published zebrafish morpholino experiment results. Homozygous HTT knockouts were embryonically viable and appeared developmentally normal through juvenile stages. Comparison of adult fish revealed significant reduction in body size and fitness in knockouts compared to hemizygotes and wildtype fish, indicating an important role for wildtype HTT in postnatal development. Our zebrafish model provides an opportunity to examine the function of wildtype HTT later in development.

## Introduction

Trinucleotide CAG repeat expansion in *Huntingtin (HTT)* is the cause of the autosomal dominant neurodegenerative disorder Huntington disease (HD) [1]. HD is characterized by progressive motor, cognitive, and psychiatric features with an age of onset inversely correlated with the length of the expanded CAG repeats. A major therapeutic approach under development for HD entails silencing of the *HTT* transcript level, although current efforts mainly focus on non-allele specific reduction of both mutant and wildtype *HTT* levels although the long-term consequences of such reductions have not been fully assessed [2].

Wildtype HTT has been shown to be important in several processes such as ciliogenesis, BDNF production, selective autophagy, and regulation of transcription [1, 3], although efforts to fully understand the effect of loss of wildtype HTT function in development have been hampered by embryonic lethality of constitutive HTT knockout mouse models [4–6]. In contrast, deletion of *htt* in *drosophila (dhtt)* appears to have little effect on viability or morphology, and dhtt appears to be dispensable for nervous system functions and development at least through juvenile stages [7]. Adult *dhtt*-knockout flies, however, exhibit accelerated neurodegeneration in a *drosophila* HD model, suggesting that HTT might carry out different functions in embryonic development and in adulthood [7].

Zebrafish has been used to model several neurodegenerative disorders due to its genetic conservation to mammals and ease of experimental manipulation [8]. In the context of HD, transgenic zebrafish expressing mutant HTT exon 1 fragment exhibit adult-onset movement deficit, which is worsened by the deletion of the N-terminal first 17 amino acids (N17) of mutant HTT fragment [9]. To understand the functions of HTT in development, several studies in zebrafish have utilized antisense morpholinos to knock down *HTT* transcripts. Such approaches have identified HTT as important in iron utilization, BDNF production, and neurulation [10–12]. Despite its usefulness, there are several drawbacks to using morpholino knockdown to study HTT functions. The short half-life of morpholino leads to knockdowns that are short-lived and would only allow insights into gene functions in early development. The inability to control the precise level of reduction in transcripts also leads to variations between morpholino experiments. Lastly, potential off-target effects are poorly characterized and often lead to gross phenotypic changes that confound the interpretation of the results [13].

Recent advances in genome editing technologies have made it possible to generate knockouts in previously understudied models. In this report, we utilized the CRISPR/Cas9 system to generate a large deletion in the genomic sequence of zebrafish *HTT (zHTT)*. Surprisingly, *zHTT* knockouts develop normally through juvenile stages with no discernible deficiencies in neurulation or gross morphological abnormalities. However, we observed a reduction in fitness in the knockout adults which is not observed in the hemizygotes, indicating that zHTT may play a more important role in adults than in early development. Our zebrafish model provides an opportunity to study the function of wildtype HTT in vertebrate development in the context of whole organisms.

## Results

### Huntingtin is well conserved between mammals and zebrafish

Full length zebrafish *Huntingtin (zHTT)* was initially sequenced and verified in 1998 [15]. Since then, 11 iterations of the genome assembly have been published covering >90% of the whole zebrafish genome [16]. In this study, we used the genome assembly from GRCz10, where the zebrafish genome contains a single copy of zHTT that encodes for a single known isoform of 3,121 amino acids (Ensembl, Fig. 1). Comparison of zHTT to human and mouse HTT reveals a high degree of conservation, with 80% similarity and ∼70% identity to both species over its entire length (Fig 1, 2A).

**Figure 1.**
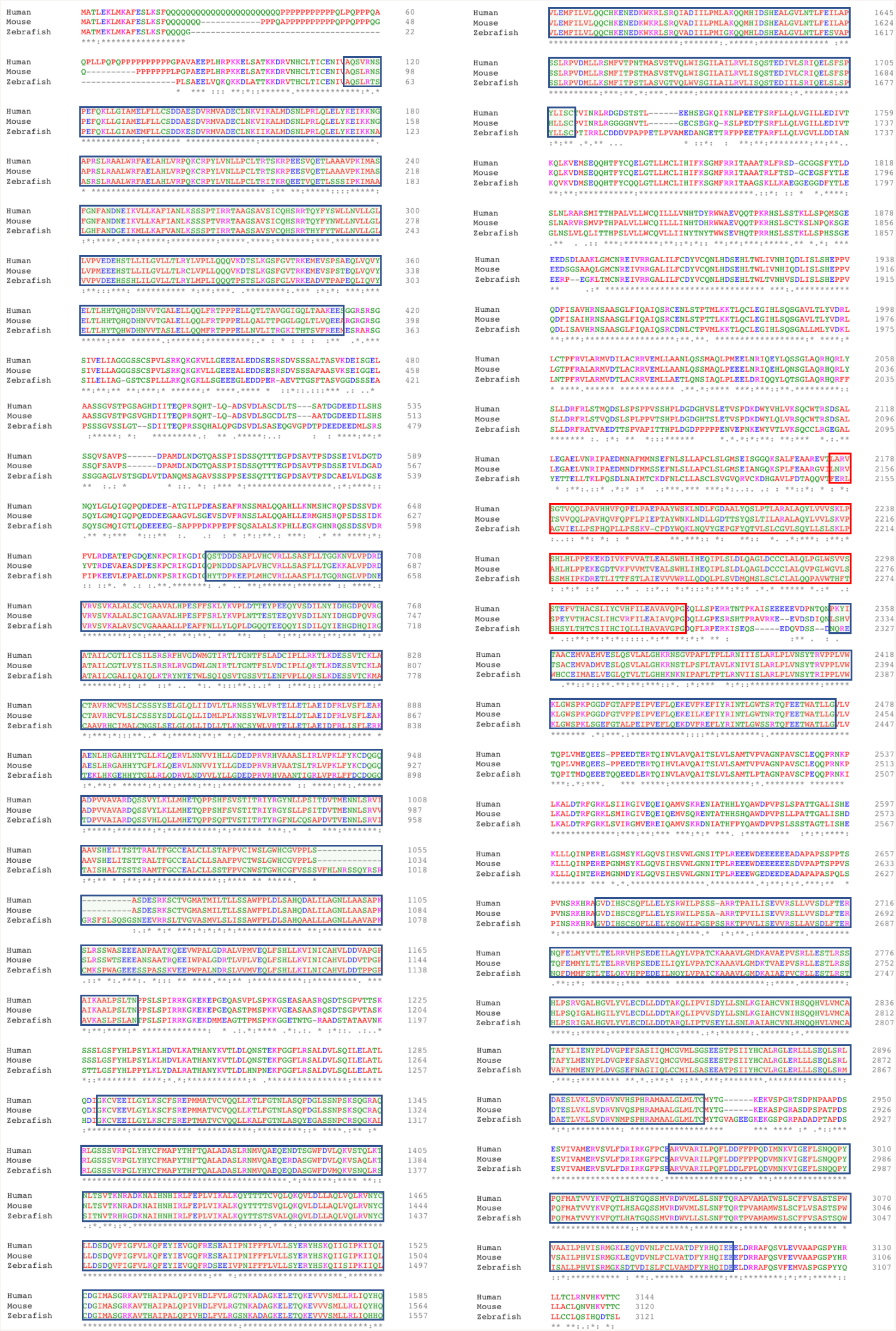
Zebrafish HTT has a high degree of conservation to mouse and human HTT. Zebrafish Amino acid sequence alignment of human, mouse, and zebrafish Huntingtin. Boxed residues indicate the locations of the HEAT domain in human. Red box indicates HEAT repeats 3 where conservation among the 3 species is the lowest.

**Figure 2.**
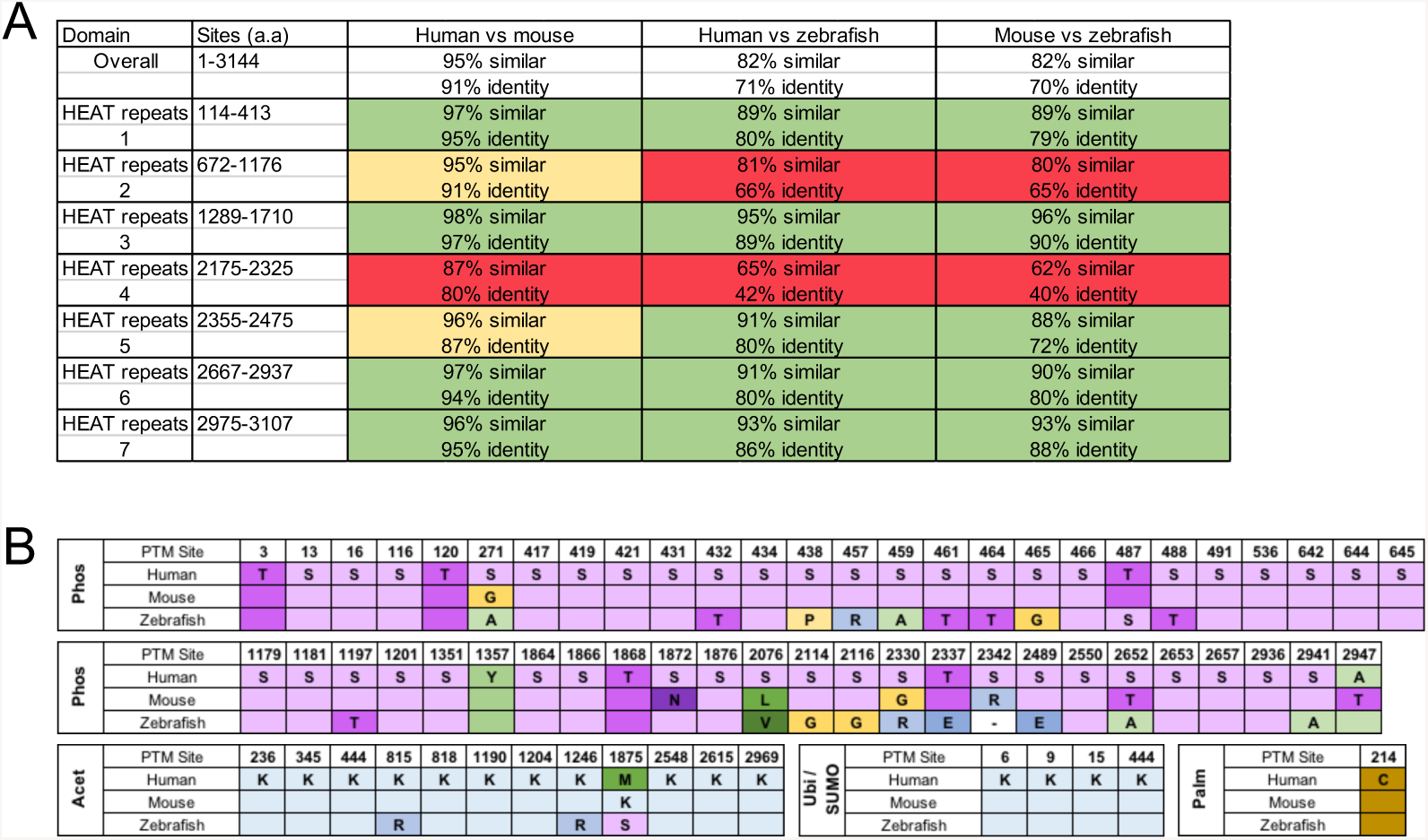
Similarity of HEAT repeat regions and post-translational modification sites between human, mouse, and zebrafish Huntingtin. (A) Comparison of amino acid similarities and identities between human, mouse, and zebrafish Huntingtin, both full-length and from different HEAT repeat stretches. Stretches that are more conserved on average are color-coded in green, while stretches that are less conserved are color-coded in red. (B) Compiled list of post-translational modification (PTM) sites and types in human HTT and their corresponding residues in mouse and zebrafish. Sites listed are obtained from [1] compiled from 10 studies, and [14]. Phos = phosphorylation; Acet = acetylation; Ubi = ubiquitination; SUMO = sumoylation; Palm = palmitoylation

To further understand the degree of conservation of HTT between humans, mouse, and zebrafish, we compared the sequences of HTT at the HEAT repeats (HTT, Elongation factor 3, protein phosphatase 2A, and TOR1) and amino acid residues which undergo post-translational modifications (PTMs), which are thought to influence HTT functions as well as modify its toxicity in the context of the HD mutation. HEAT repeats have been proposed as potential scaffold sites for HTT-interacting proteins [1] and we hypothesized that higher similarities in these domains in vertebrate HTT might indicate functional conservation. At the amino acid sequence levels, we observed that HEAT repeats 1, 3, 5, 6, and 7 have higher than average conservation, indicating that these stretches potentially carry important functions of the protein across species (Fig. 2A). HTT also undergoes a large number of PTMs, although the role of these PTMs have mostly been studied in the context of mutant HTT. We summarized these PTMs, compiled from [1, 14], and compared their conservation across the three species in (Fig. 2B). Overall, 60 out of 68 (88.2%) identified PTM residues are conserved between human and mouse, compared with 45 (66.2%) in zebrafish (Fig. 2B). Overall, we conclude that HTT is highly conserved with high levels of sequence identity and similarity between human HTT and zHTT, including protein domains and PTM residues of importance to HTT function.

### CRISPR/Cas9 deletion of genomic zHTT

The genomic locus of *zHTT* contains 67 exons spanning 80kbp. To inactivate zHTT, we used CRISPR/Cas9-mediated genome targeting with 2 guide RNAs (gRNAs) targeting exon 1 and exon 7 (Fig. 3A). Following injection of Cas9 and both gRNAs into 1-cell stage embryos, we screened for successful deletion of the genomic region flanked by both gRNAs using distally located primers (Fig. 3B). These primers flanked a region that is ∼11kbp in wildtype fish, but is reduced to <700bp upon successful deletion. Sanger sequencing of PCR product from injected zebrafish confirmed that the deletion successfully removed the genomic region spanning exon 1 and intron 7 of the *zHTT* locus (Fig. 3C).

**Figure 3.**
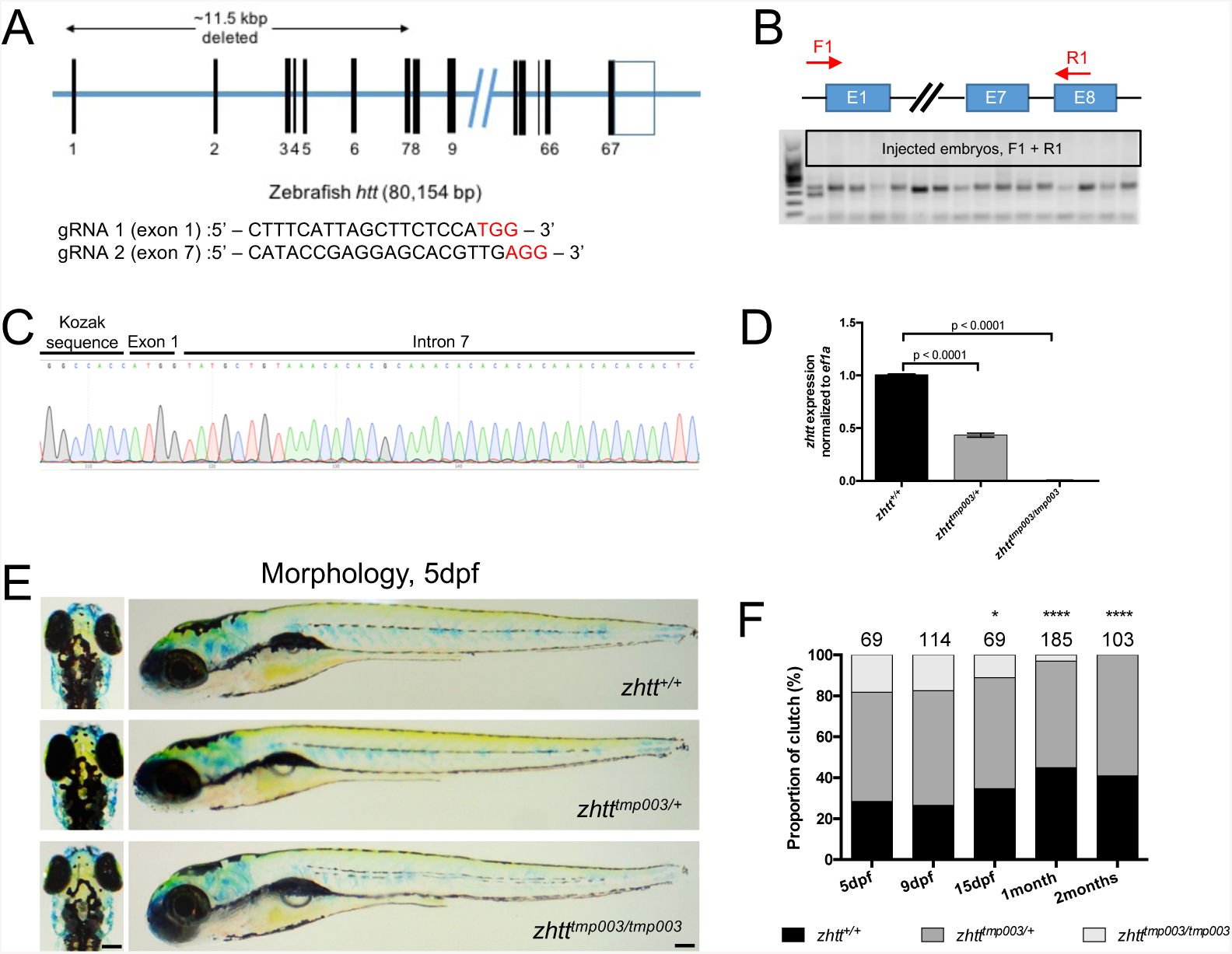
Generation and characterization of HTT hemizygous and knockout zebrafish. (A) Graphical representation of *zhtt* genomic locus and sequences of sgRNAs used for CRISPR/Cas9 editing. (B) Distal primers F1/R1 are able to detect the presence of large genomic deletion on injected embryos. (C) Representative Sanger sequencing trace of injected embryo showing that large deletion removes a large portion of Exon 1 to Intron 7 of *zhtt*. (D) qRT-PCR analysis of *zhtt* mRNA expression levels of 5 dpf wildtype, hemizygous, and knockout zHTT embryos. Mean values ± SEM. n = 2 pools of 12 zebrafish each at 72 hpf for each genotype. p-values determined by one way ANOVA with Dunnett’s post hoc test. (E) Morphology of 5 dpf zhtt wildtype, hemizygous, and knockout zebrafish. Lateral and dorsal views. Scale bar = 100µm. (F) Genotypic ratios of progenies from heterozygous intercrosses of *zhtt*^*tmp003/*+^. Dotted lines represent expected ratio of each genotype. Number of fish analyzed per clutch is represented above the bars. *zhtt*^*tmp003/tmp003*^ are significantly below expected ratio at 15dpf, 1 month, and 2 months. p-values determined by Chi-square analysis. *p = 0.0265; ****p < 0.0001. dpf, days postfertilization.

We raised a portion of injected zebrafish embryos to adulthood and screened for adults that transmit this large deletion into the next generation. We then identified and raised adult zebrafish that are heterozygous for the large deletion, subsequently referred to as the *zhtt*^*tmp003*^ allele. Heterozygous intercrosses generated embryos of all three genotypes in expected ratio. We confirmed that the deletion leads to loss of *zHTT* expression by qPCR of 3 days postfertilization (dpf) embryos (N = 20 embryos/genotype) using primers that recognize sequences immediately downstream of the deletion site (Fig. 3D). Taken together, we have successfully established a zebrafish knockout line for zHTT with a large, heritable genomic deletion.

### zHTT hemizygous and knockout embryos do not exhibit gross morphological defects

Multiple morpholino experiments to knock down zHTT have suggested an important role for zHTT in early development [10, 11]. Morpholino-induced *zHTT* knockdown (zHTT-MO) zebrafish embryos display smaller yolk extension, smaller head and eye size, as well as curvature of the body axis. Defects were shown to manifest starting at 24 hours postfertilization (hpf) and remain visible up to 5 dpf. Strikingly, we do not see gross morphological phenotypes in our hemizygous and homozygous zHTT knockout embryos at 5dpf (Fig. 3E, N > 100 embryos/genotype). At these stages, *zhtt*^*tmp003/*+^ and *zhtt*^*tmp003/tmp003*^ embryos are morphologically indistinguishable from their wildtype siblings. These results indicate that there are fundamental differences between morpholino knockdown and genomic knockout of zHTT.

### zHTT knockouts, but not hemizygous, show reduced size and fitness in adulthood

One of the disadvantage of morpholinos is their short half-life, leading to a small window of time in which they are effective in knocking down transcript levels. In contrast, genetic zHTT knockouts and hemizygotes provide an opportunity to study the consequences of HTT deficiency in vertebrates at more advanced developmental time points. To this end, we raised progenies from heterozygous intercrosses and sampled them at various time points in development. We observed that while zHTT hemizygotes and knockouts are initially obtained in Mendelian ratios, zHTT knockouts have a decreased fitness and survive below expected proportions starting at 15dpf and more significantly by 1 month of age (Fig. 3F, N > 60 per time point, Chi-square test, p < 0.0001).

### zHTT hemizygous and knockout embryos do not exhibit defects in neurulation and iron utilization

Morpholino-mediated knockdown of zHTT has been shown to result in abnormalities in neural tube morphogenesis that is quantifiable at 24 hpf [12], as well as deficiency in iron utilization that manifests as blood hypochromia in early embryos [10]. To examine whether these phenotypes contribute to reduced fitness in zHTT knockouts, we imaged the heart regions of embryos obtained from heterozygous *zhtt*^*tmp003/*+^ intercrosses (Fig. 4A). In contrast to zHTT-morpholino studies, there is no apparent hypochromia phenotype that accompanies the decrease in zHTT levels in either *zhtt*^*tmp003/*+^ or *zhtt*^*tmp003/tmp003*^ embryos at 2 dpf.

**Figure 4.**
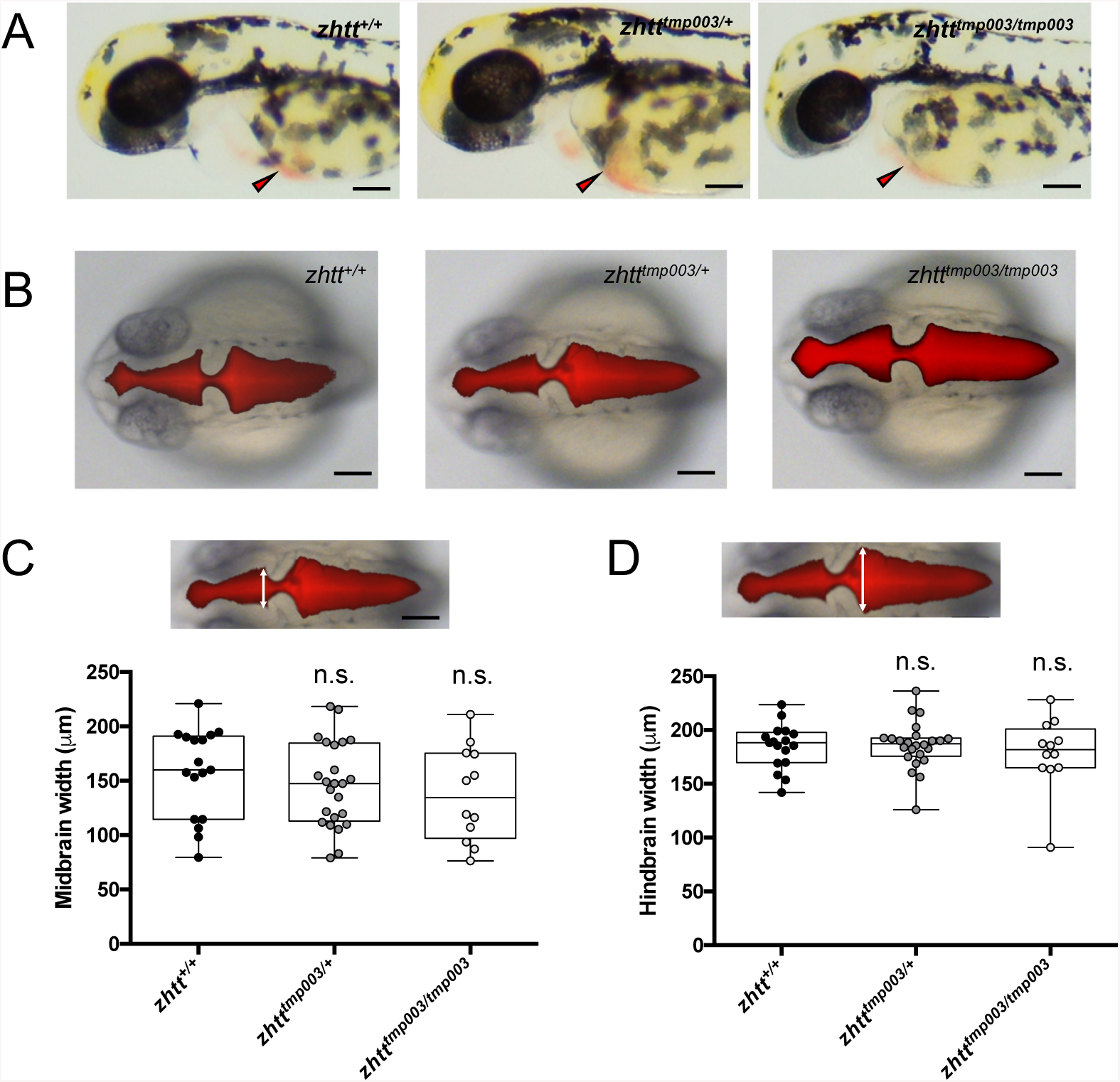
zhtt-deficient zebrafish display normal iron utilization and neurulation early in development. (A) Representative images of 2dpf wildtype, hemizygous, and knockout zHTT embryos. Red arrowheads point to the hearts of each embryos. No apparent hypochromia observed. Scale bar = 100µm. (B) Representative images of 24hpf Texas Red-Dextran injected wildtype, hemizygous, and knockout zHTT embryos. Scale bar = 100µm. Measurement of midbrain (C) and hindrain (D) widths show no significant difference in neurulation observed between the genotypes. Statistical significance calculated using one way ANOVA with multiple comparisons between groups analysed by Dunnett’s test.

To visualize potential defects in neurulation in the midbrain and hindbrain regions of zHTT-deficient embryos, we injected Texas Red-labeled dextran into the hindbrain of 24 hpf embryos, followed by imaging and measurement (Fig. 4B, C). Reduction in *zHTT* transcript has previously been reported to alter neuroepithelial cell formation resulting in constriction along the neural tube that is attributed to the interaction between HTT and ADAM10-NCadherin [12]. We did not find significant differences in the sizes of midbrain or hindbrain across all genotypes at 24 hpf (Fig. 4B-D). Taken together, these results suggest that reduction in zHTT knockout survival is not a result of early embryonic defects in neurulation or iron utilization.

### zHTT knockout adults are smaller compared to wildtype siblings

Although embryonic development appears unaffected in zHTT knockouts and hemizygotes, we observed a reduction in body size in adulthood (30dpf) specific to zHTT knockouts but not hemizygotes (Fig. 5A, B) (wildtype vs knockouts, p=0.0040, one way ANOVA with Dunnett’s multiple comparisons). We hypothesized that competition between fish of different genotypes might contribute to this size difference potentially leading to the genotype-specific lethality observed in zHTT knockouts. To test whether this reduction in survival reflects intrinsic deficits linked to the levels of zHTT or is affected by other extrinsic factors such as competition, we genotyped progenies of *zhtt*^*tmp003/*+^ heterozygous intercrosses at 5 dpf and grew each genotype separately out of competition from other genotypes (Fig. 5C). When compared to their wildtype siblings (N = 36), neither conditions improved the survival of zHTT knockouts (Fig. 5C, p < 0.0001). In addition, we observed that zHTT knockouts grown out of competition do not survive longer or at higher proportions compared to zHTT knockouts grown in competition with siblings of other genotypes (N = 21 knockouts grown out of competition). Taken together, these observations suggest that there is an intrinsic requirement for HTT essential for maintaining viability in adulthood.

**Figure 5.**
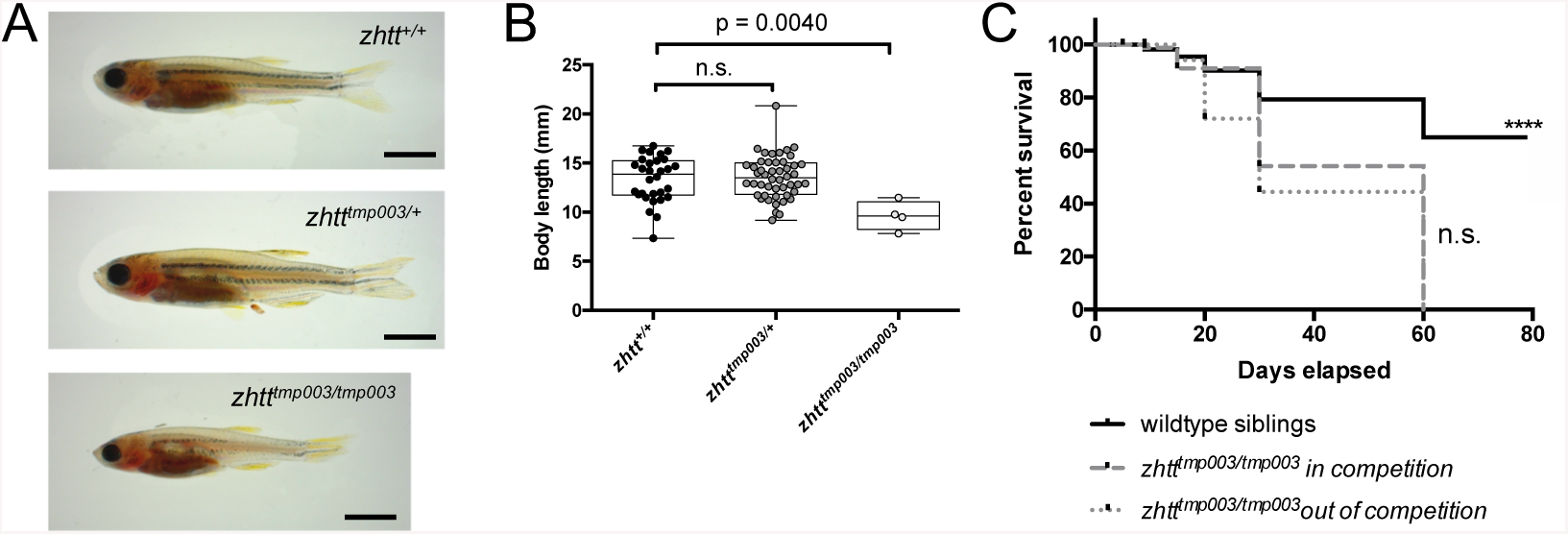
zHTT-knockout, but not hemizygous, exhibit reduced size and survival. (A) Representative images of 30dpf wildtype, hemizygous, and knockout zHTT adults. Scale bar = 2mm. (B) Body length measurements of wildtype (N = 30), hemizygous (N = 48), and knockout (N = 4). Statistical significance calculated using one way ANOVA with multiple comparisons between groups analysed by Dunnett’s test. Significant difference in body lengths was observed between wildtype and knockout groups. (p=0.0040). (C) Survival curves of zHTT knockout embryos when raised in competition with fish of other genoypes or out of competition starting at 5dpf. No significant difference in the lengths of survival observed in zHTT knockouts between the two treatments. Both conditions show reduced survival when compared to wildtype siblings (p <0.0001).

## Discussion

In this study, we described a new zebrafish model to understand the function of HTT in the neonatal period in vertebrates through genetic deletion of HTT. We showed that while HTT confers fitness and viability in adult zebrafish, its deletion does not lead to embryonic lethality as previously thought. Our findings are in line with previous studies which showed that embryonic phenotypes in rodent knockout models for HTT can be rescued by HTT expression in the extraembryonic tissues, suggesting that while HTT is essential in mammalian development, the essentiality of its function during development largely related to extra-embryonic, non-zygotic tissues [17]. This is further supported by the lack of developmental phenotypes in *drosophila htt* knockout models, although the low conservation between vertebrate and invertebrate HTT implies that there might be other mechanisms at play.

The differences between our findings and previous reports can be attributed to several factors. Firstly, recent publications have substantiated the possibility that genetic knockouts of deleterious genes are often compensated and tolerated by the upregulation of other genes in the same pathways, masking the phenotypes that are otherwise obtained with knockdown experiments [18]. The lack of full understanding of HTT functions in vertebrates makes it hard to predict if other genes with the potential to compensate for the loss of HTT function exists in zebrafish. Secondly, it is possible that at least some of the phenotypes described in *zHTT*-morpholino experiments reflect off-target effects. Indeed, morphological changes such as curved body axes, pericardial oedema, as well as reduction in head and eye sizes are part of the spectrum of non-specific morpholino effects [19]. In addition, several of these effects are also dose-dependent, possibly leading to a wide range of phenotypes depending on the level of knockdown achieved.

Overall, the zebrafish HTT knockout model we describe provides an opportunity to study the function of HTT during the neonatal period in the context of the whole organism.

## Materials and Methods

### Ethics statement

All zebrafish experiments were approved by the Singapore National Advisory Committee on Laboratory Animal Research.

### Zebrafish maintenance and husbandry

Zebrafish were maintained in Biological Resources Center Zebrafish facility (A*STAR, Singapore) according to Institutional Animal Care and Use Committee guidelines (IACUC #151094). Zebrafish were reared at 28.5°C. Embryos were obtained from natural mating and kept in egg water up until 5dpf. Unless otherwise stated, all zebrafish lines were from the AB wildtype background.

### CRISPR/Cas9 deletion

nlsCas9 protein was produced by NTU Protein Production Platform (www.proteins.sg). Single guide RNAs (sgRNAs) were made following protocol from [20]. Primers used for sgRNA construction are as follows: gRNA 1: 5’-TTAATACG ACTCACTATAGGCAAGGCCCGCTGTCGGCCGAGGGTTTTAGAGCTAGAAATAGC; gRNA 2: 5’-TTAATACGACTCACTATAGGCATACCGAGGAGCACGTTGAGGGTTTTAGAGCTAGAAATAGC.

### Genotyping

Genomic DNA was obtained either from whole embryos, plucked scales from adult fish, or from tail fin biopsy of 3 dpf zebrafish (following methods from [21]). Tissues were lysed by incubation in 10-20 µl of 25mM NaOH / 0.2mM EDTA at 98°C for 10 minutes, followed by neutralization using equal volume of 40mM Tris-HCl pH 8.0. Multiplex PCR was performed using the primers zf-mp-lp 1 (5’-AGGCTGACTATTGTTGTTGTTG-3’),zf-mp-rp 1 (5’-CAAGTAGCGCAGTGTCAGTAAT-3’), and zf-mp-rp 2 (5’-TTGAGTGTGTG TTTGAGCTGTA-3’). PCR was performed using Taq DNA polymerase (Invitrogen) with the following conditions: 94°C (2 mins), 35 cycles of 94°C (30 sec) 56°C (30 sec) 72°C (45 sec), 72°C (5 mins). Expected PCR band lengths are 503bp and 327bp, corresponding to the deleted and wildtype zHTT alleles respectively.

### RNA isolation, cDNA synthesis, and qPCR

5 dpf embryos from heterozygous inter-crosses were subjected to tail fin biopsies. Tail fin pieces are used for genotyping while trunk and head fragments are frozen down in RNA extraction buffer (Favorgen) and stored in −80°C until genotypes of each embryo is known. 10-12 embryos per genotype were pooled for RNA extraction using the Total Tissue RNA Mini Kit (Favorgen) and total RNA is eluted in 15 µl volumes. 1-2 µg of total RNA for each genotype was converted to cDNA using the High Capacity cDNA Reverse Transcription Kit (Thermo Fisher). qPCR was performed using SYBR Select Master Mix (Thermo Fisher). qPCR primers are as follows: ef1a − 5’-AGGACATCCGTCGTGGTAAT-3’ and 5’-AGAGATCTGACCAGGGTGGTT-3’ zhtt − 5’-atcattccagcagcagcaag-3’ and 5’-tttctggaactctggagacg-3’.

### Brain ventricle microinjection

Injection of Texas-Red Dextran (Thermo Fisher) was performed following protocol described in [22]. In vivo fluorescent images were obtained with an upright laser scanning microscope (LSM) Meta 510 (Carl Zeiss) equipped with an AxioCam HRC digital camera (Carl Zeiss Vision) and 543nm laser line. Brightfield and fluorescent images were combined using Photoshop (Adobe).

### Imaging

Brightfield images of embryonic and adult zebrafish were taken on an Olympus SZ61 Stereo Microscope with Moticam 580 camera attachment.

### Statistics

All statistics analyses were performed in Prism 6 (Graphpad).

## Conflict of interest

The authors declare no conflict of interest.

## Author Contributions

H.S. and M.A.P. conceived and planned the study. H.S. and C.J.A. performed experiments. H.S. and C.J.A. analyzed data. H.S. and M.A.P. wrote and revised the manuscript.

## Acknowledgments

We thank the staff of the Biological Resources Center zebrafish facility (Agency for Science, Technology, and Research, Singapore). M.A.P. is supported by grants from the Agency for Science, Technology and Research, and the National University of Singapore, Singapore.

## Notes

#### Summary of Updates

Author order has been revised

